# Neurodegeneration in the olfactory system in Niemann Pick type C1 disease

**DOI:** 10.1101/2025.11.22.689959

**Authors:** Maria Grazia Fioriello, Donatella Lobraico, Raffaella Pia Gatta, Simona Lobasso, Johannes Reisert, Antonio Frigeri, Michele Dibattista

**Affiliations:** Department of Translational Biomedicine and Neuroscience (DiBraiN), University of Bari Aldo Moro, Bari, Italy; Monell Chemical Senses Center, Philadelphia, PA, United States

## Abstract

*Niemann-Pick type C1 (NPC1) disease* is a rare neurodegenerative disorder linked to defective cholesterol biosynthesis and altered lipid regulation. In patients, the most common variant in the *Npc1* gene is a missense mutation that leads to a misfolded and non-functional NPC1 protein. We used a mouse model carrying this mutation (I1061T, *Npc1*^tm(I1061T)Dso^) to investigate the olfactory system since olfaction is often negatively impacted in many neurodegenerative disorders, with olfactory decline frequently preceding other neurodegenerative symptoms.

We characterized cell types in the olfactory epithelium (OE) in wild-type, heterozygous, and homozygous *Npc1*^tm(I1061T)Dso^ mice to analyze neurodegeneration at two time points, namely 36 and 60 days after birth. In homozygous *Npc1*^tm(I1061T)Dso^ mice the density of olfactory sensory neurons (OSNs) at 36 days after birth is decreased compared to wild type and heterozygous mice while the density of immature OSNs and the stem cell niche seems to not be affected. In the OE of the homozygous *Npc1*^tm(I1061T)Dso^ mice we found an increased density of apoptotic cells and clear sign of neuroinflammation in macrophage/microglia infiltrating the OE. We analyzed the lipid profile of OE by positive MALDI-TOF and found increased markers of neuroinflammation homozygous *Npc1*^tm(I1061T)Dso^ mice.

These structural changes affected the functionality of the OE since we found reduced odorant responses in *Npc1*^tm(I1061T)Dso^ mice compared to wild type. Additionally, our olfactory behavioral tests revealed deficits in odor-guided food-seeking tests in *Npc1*^tm(I1061T)Dso^ but not in the wild type and heterozygous mice. We also analyzed the olfactory abilities of a family of three with the parents carrying two different mutations of the *Npc1* gene, with the child carrying both mutations and being diagnosed with NPC1 disease. Using the Sniffin’ Sticks olfactory test, we found that all three were hyposmic with the child having severe hyposmia.

Our work extensively characterized the OE structurally and functionally proposing it as a sentinel to monitor disease progression. We are the first to show that NPC1 patients are affected by severe hyposmia and that heterozygous parents could be monitored for olfactory abilities since it could be a biomarker for future neurological disorders.

## Introduction

Niemann-Pick type C (NPC) disease is an autosomal recessive neurodegenerative disease caused by mutations in the *Npc1* gene (in 95% of patients) or NPC2 gene (Pentchev 2004). Both genes encode proteins required for cholesterol export from lysosomes (Rosenbaum and Maxfield 2011). The NPC1 glycoprotein is involved in the efflux of low density lipoprotein (LDL) derived unesterified cholesterol from late endosomes and lysosomes to other intracellular compartments (Sokol et al. 1988; Liscum et al. 1989). The NPC2 protein binds cholesterol (Friedland et al. 2003) and mutations in NPC2 produce accumulations of unesterified cholesterol, sphingolipids and complex gangliosides in late endosomes and lysosomes of most cell types of the body indistinguishable from those of NPC1 mutations (Pacheco and Lieberman 2008).

Loss-of-function mutations in either of these proteins lead to deficient lipid trafficking, abnormal regulation of cholesterol biosynthesis, and intracellular accumulation of unesterified cholesterol and gangliosides GM2 and GM3 (sialic acid-containing sphingolipids, major constituents of neuronal membranes) in late endosomes/lysosomes (Hovakimyan et al. 2013, Vasques et al. 2023), due to impaired mobilization and re-esterification of LDL cholesterol throughout the body, especially in liver, spleen, lung, and brain (Neufeld et al. 1999).

The most prevalent mutation in the *Npc1* gene, I1061T, accounts for approximately 15–20% of all disease-associated alleles in humans (Millat et al. 1999; Davies and Ioannou 2000; Park et al. 2003). This variant leads to the misfolding of the NPC1 protein, which is subsequently directed toward ER-associated degradation (Gelsthorpe et al. 2008). This mutation predisposes patients to the classic NPC1 disease clinical phenotype (Yamamoto et al. 2000; Millat et al. 1999), which includes hepatosplenomegaly, ataxia, dystonia, and progressive neurodegeneration (Garver et al. 2007) (in late-infantile and juvenile forms between 3 and 15 years of life, 60–70% of cases) (Pacheco and Lieberman 2008).

Neurodegeneration can even appear later with onset in adolescence and adulthood when it is characterized by cognitive, auditory and coordination impairments (Geberhiwot et al. 2018). Animal models for NPC1 disease reported age-dependent neuronal loss in the prefrontal cortex, thalamus, and brainstem, as well as degeneration of cerebellar Purkinje cells, astrogliosis, upregulation of inflammatory genes, and accumulation of cholesterol-derived compounds and sphingolipids within hepatic and neural tissues. These changes closely parallel those described in human NPC1 disease (Vanier 1983; 1999; Walkley and Suzuki 2004; Cologna et al. 2014).

Except for retinal degeneration (Mauch et al. 2001), sensory systems such as olfactory, trigeminal, or auditory pathways in NPC1 disease have not been studied so far. Neurodegenerative diseases e.g. Parkinson’s disease show early olfactory impairment, which occurs at least two years before motor symptoms become evident (Berendse et al. 2001; Attems and Jellinger 2006; Doty 2012). The same applies to Alzheimer’s disease (Attems and Jellinger 2006, 2010; Jellinger and Attems 2010; Wesson et al. 2010; Bahar-Fuchs et al. 2011; Dibattista et al. 2020). The most peripheral part of the olfactory system is the olfactory epithelium (OE) located in the upper nasal cavity, and it consists of several main cell types: the olfactory sensory neurons (OSNs), the sustentacular cells (SUSs) and two populations of reserve stem cells located in the basal lamina of the OE, named horizontal basal cells (HBCs) and globose basal cells (GBCs). OSNs are bipolar neurons responsible for the detection of odorant molecules that enter the nasal cavity via breathing or sniffing. SUSs maintain the microenvironment in the OE regulating OSNs function and basal cells proliferation (Lobraico et al. 2025). Basal cells in the OE support neurogenesis that leads to the formation of new functional OSNs throughout life and supports a continuous turnover of these cells. This turnover is very remarkable and is unique in the nervous system of mammals ((Roskams et al. 1998; Nickell et al. 2012; Liang 2020). Both types of basal cell populations are multipotent, with HBCs considered reserve stem cells, whereas GBCs are responsible for continuous replacement of OSNs and other cell types in the OE. (Graziadei and Graziadei 1979; Schwob et al. 2017). Because of these unique features, we hypothesized that the olfactory system and in particular the OE could be exploited to better understand the ongoing maintenance of a sensory system and the potentially ensuing neurodegeneration. Therefore, we studied whether structural and functional changes in the OE of a mouse model carrying the *Npc1*^I1061T^ mutation occur and what their time course might be in similarity to other neurodegenerative diseases where olfactory decline precedes other symptoms. We found that the cell composition and functionality of the OE are impaired in *Npc1*^I1061T^ mutant mice as early as post-natal day 36. In particular, we found that the density of mature OSNs as well as the odorant response decreased in *Npc1*^I1061T^ mutant mice (MUT) compared to heterozygous (HET) and wild type (WT) mice. We found infiltrated macrophages in the OE and increased number of apoptotic cells. In addition, we found that even HET mice showed altered features similar to MUT. In addition, we could test two parents with heterozygous mutations of the *Npc1* gene and their young child carrying both mutations. We found that all three, although to varying degrees, had impaired olfactory function.

## 1. Results

### NPC1 mutant mice exhibit early neurodegeneration

In NPC1 disease impairment of lipid trafficking affects the CNS resulting in neuron loss throughout the brain (Walkley and Suzuki 2004). We hypothesized that in mice homozygous for the Npc1 I1061T mutation (MUT), neurodegeneration could also be observed in the olfactory epithelium (OE), an outpost of the brain located in the nasal cavity. We performed immunofluorescence on OE slices at 36 days after birth (P36) and at a late time point P60 (Fig. 1) by using an antibody against olfactory marker protein (OMP), a marker of mature OSNs (Margolis 1972; Hartman and Margolis 1975; Margolis 1982; Kondo et al. 2010; McClintock et al. 2020). We found that the density of OMP-positive neurons decreases in mutant mice (Fig. 1h, i) compared to WT (Fig. 1b, c) and HET (Fig. 1e, f). Since the continuous proliferation and generation of new OSNs in the OE, some OSNs are not yet mature (aka immature) and we used a mouse anti β-III tubulin (Tuj1) antibody as a marker for immature OSNs (Gavid et al. 2023). We could clearly see Tuj1+/OMP- neurons just above the basal layer of the OE. A third cell population was represented by neurons that were showing some degree of colocalization between Tuj1 and OMP. We considered cells immature only Tuj1 positive and OMP negative (Tuj1+/OMP-, Fig. 1a, d, g, j, m, p). Immature cells were not significantly different between the three genotypes at P36 (Fig. 1a, d, g, summary plot in s). In addition, when Tuj1+/OMP+ cells were quantified, we could observe a decreased density in MUT compared to HET and WT. Interestingly, although the overall neuronal cell population decreases (see Tuj1+/OMP+ and OMP+ in Fig. 1 and summary in Fig. 1 s), immature cells seem to remain constant in their density across genotypes. The same staining was performed at P60 and found the same patterns: a decrease in OMP+ density in the MUT mice (Fig. 1 j-r and summary in t) and no change in Tuj1+/OMP- cell density across genotypes. At P60 we noticed the OMP signal was restricted to a more apical region of the OE (closer to the ciliary layer) in MUT mice (Fig. 1q).

**Figure 1.**
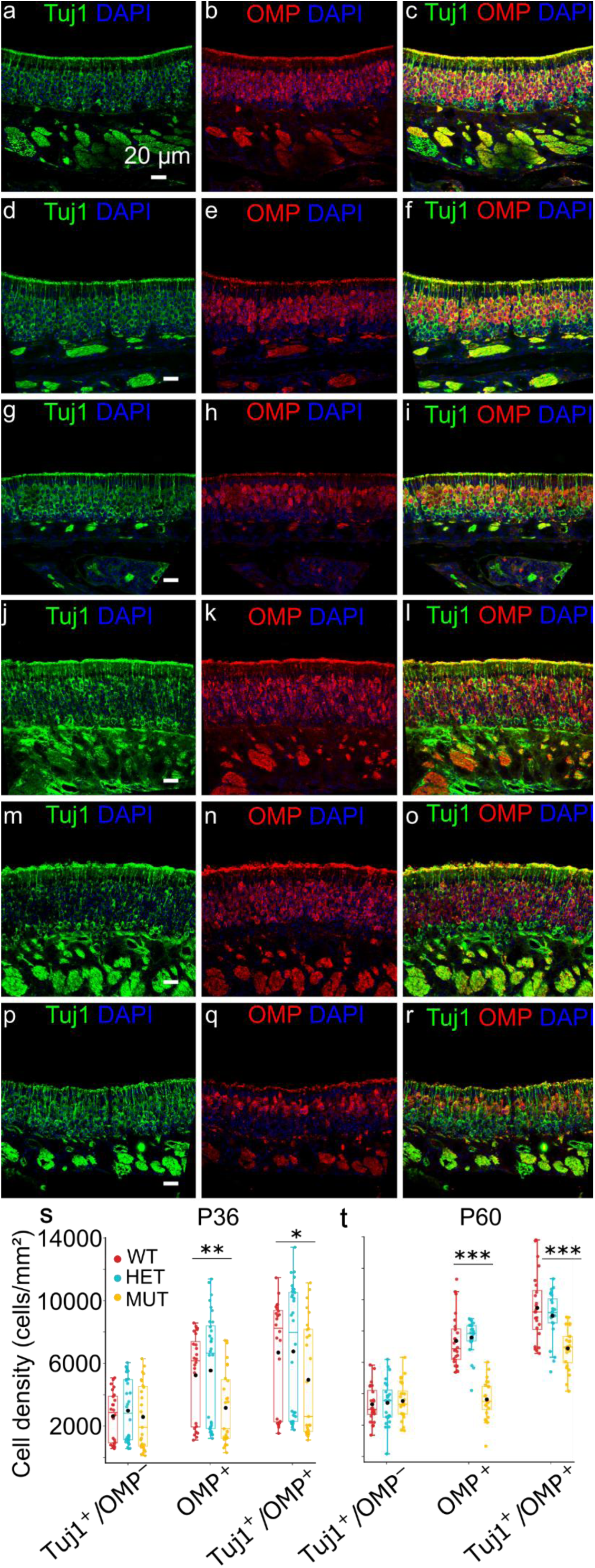
OSNs early neurodegeneration in OE. Mature OSNs and total OSN cell density (cells/mm2) decrease in MUT mice at P36 and at P60. Coronal sections from WT (a, b, c, j, k, l), HET (d, e, f, m, n, o) and MUT (g, h, I, p, q, r) olfactory epithelium showing from the left to the right green Tuj1+ (immature and transitory OSNs), red OMP+ (mature OSNs), and orange staining Tuj1+/OMP+ in mice ages P36 and P60. (s) Box plot showing cell density in WT, HET and MUT mice at P36 and at P60 (t). The central black point represents the mean, central line: median, upper and lower box boundaries: 25th and 75th percentile, extreme lines: The highest and lowest value. (Kruskal–Wallis test followed by Mann-Whitney and Benjamini–Hochberg (BH) post hoc analysis were performed for OMP+: H(2) = 12.77, p = 1.69e-3 and Tuj1+: H(2) = 8.75, p = 1.26e-2; p = *0.05, **0.01, *0.001. Sample size: n = 3 mice for each genotype at P36.) (t) Kruskal–Wallis test followed by Mann-Whitney and Benjamini–Hochberg (BH) post hoc analysis was performed to OMP+: H(2) = 45.43, p = 1.36e-10 and one-way ANOVA, followed by Tukey test post hoc analysis was performed to Tuj1+ (genotype: F(2, 69) = 15.79, p = 2.25e-6): p = *0.05, **0.01, ***0.001. Sample size: n = 3 mice for each genotype at P60. For more detailed on the post-hoc analysis refer to the table 1.

### OSNs undergo apoptosis during neurodegeneration

Apoptosis is an hallmark of many neurodegenerative diseases, and we used cleaved Caspase3 (cCasp3), a marker for activated caspase-3 (Snigdha et al. 2012; Ponder and Boise 2019) to test whether apoptotic cells are present in the OE. Indeed, apoptotic cells increased at P36 in MUT mice (Fig. 2c, summary in g) compared to HET (Fig. 2b) and WT (Fig. 2a). At P60 apoptosis is already enhanced in HET (Fig. 2e) and MUT mice compared to WT (Fig. 2d, summary in g bottom panel). These results may explain the decreased density of OMP+ cell types that we observed in the OE of MUT mice.

**Figure 2.**
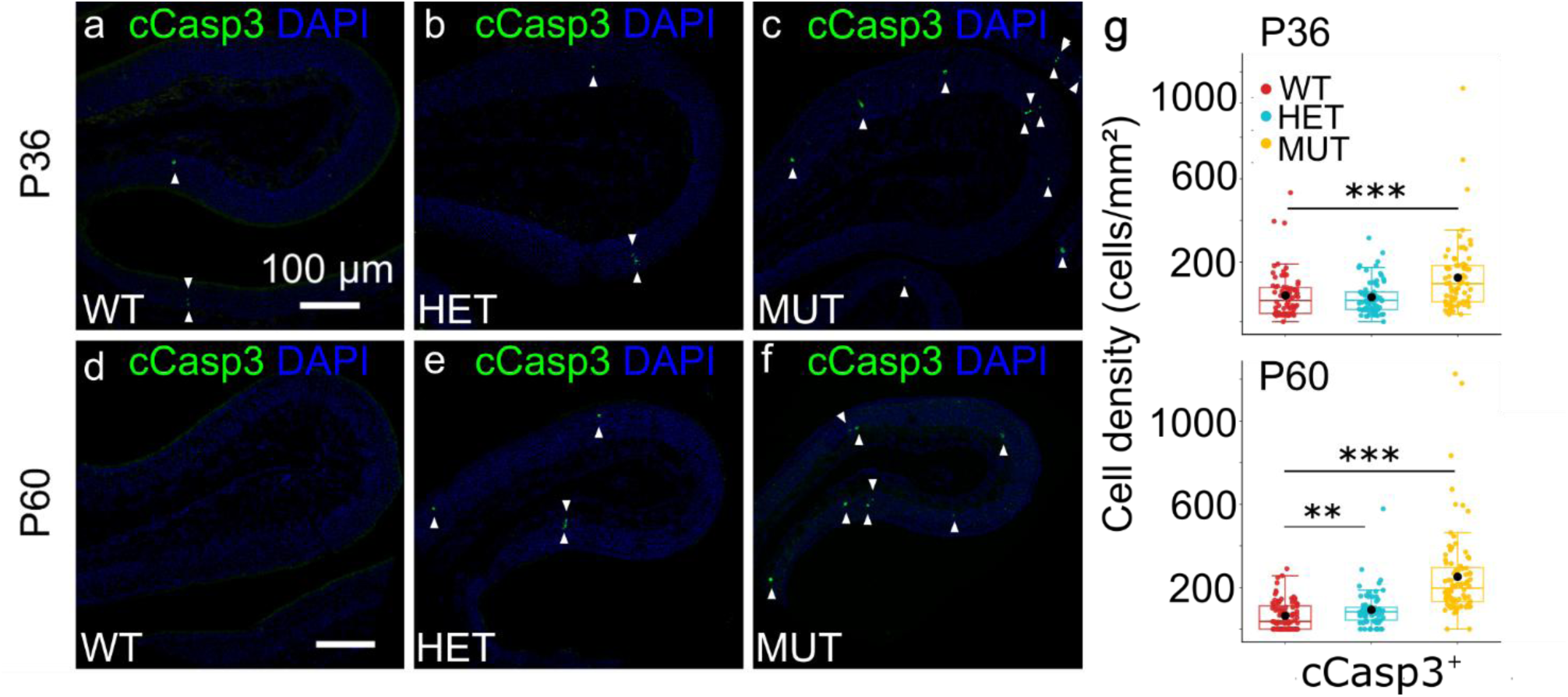
Increased apoptotic cell density in MUT mice. (a – f) Coronal sections from WT, HET and MUT olfactory epithelium showing apoptotic cells (cCasp3+) in mice ages P36 and P60. White arrowheads indicate cleaved Caspase III apoptotic neurons in OE. (g) Box plots showing cell density in WT, HET and MUT mice at P36 in the upper panel and P60 in the lower panel. The central black point represents the mean, the central line: median, upper and lower box boundaries: 25th and 75th percentile, extreme lines: the highest and lowest value. Kruskal–Wallis test followed by Mann-Whitney and Benjamini–Hochberg (BH) post hoc analysis was performed to cCasp3+: H(2) = 23.58, p = 7.59e−6 at P36 and at P60: H(2) = 103.39, p = 3.53e−23; **0.01, ***0.001. Sample size: n = 3 for each genotype and for each age. For more detailed on the post-hoc analysis refer to the table 1.

### NPC1 mutant mice exhibits neuroinflammation in OE

What does trigger apoptosis in the OE of MUT mice? We sought to investigate microgliosis since it is a common finding in NPC1 disease models (Hovakimyan et al. 2013; Praggastis et al. 2015a; Seo et al. 2016). We monitored microglia/macrophage activation by using antibodies against Iba1, the ionized calcium-binding adapter molecule 1 a widely recognized microglia/macrophage marker (Ito et al. 1998).

Microglia/macrophages recruitment is more visible in MUT (Fig. 3c, i, f, l) and HET (Fig. 3b, h, e, k) mice compared to WT (Fig. 3a, g, d, j) as early as P36 (Fig. 3 c, i). While Iba1^+^ cells in WT are mostly localized below the basal lamina or lying over it (Fig. 3 a, g, d, j), in HET (Fig. 3 b, h, e, k) and MUT (Fig. 3 c, i, f, l) they penetrate the epithelium. Both at lower and higher magnification we compared the distribution of Iba1+ cells in the OE of HET (Fig. 3 b, h, e, k) and MUT (Fig. 3 c, i, f, l) with WT. In addition, the activation of macrophages in MUT mice (Fig. 3 i, l) was confirmed by their morphology that became amoeboid and hypertrophic, characterized by large cell bodies with shorter thicker processes (Fig. 3 m-r). In HET mice it seems that macrophages had a more intermediate phenotype (Fig. 3 n, q). In summary, a decrease of OMP+ positive OSNs may be triggered/exacerbated by neuroinflammatory processes in the OE of MUT mice.

**Figure 3.**
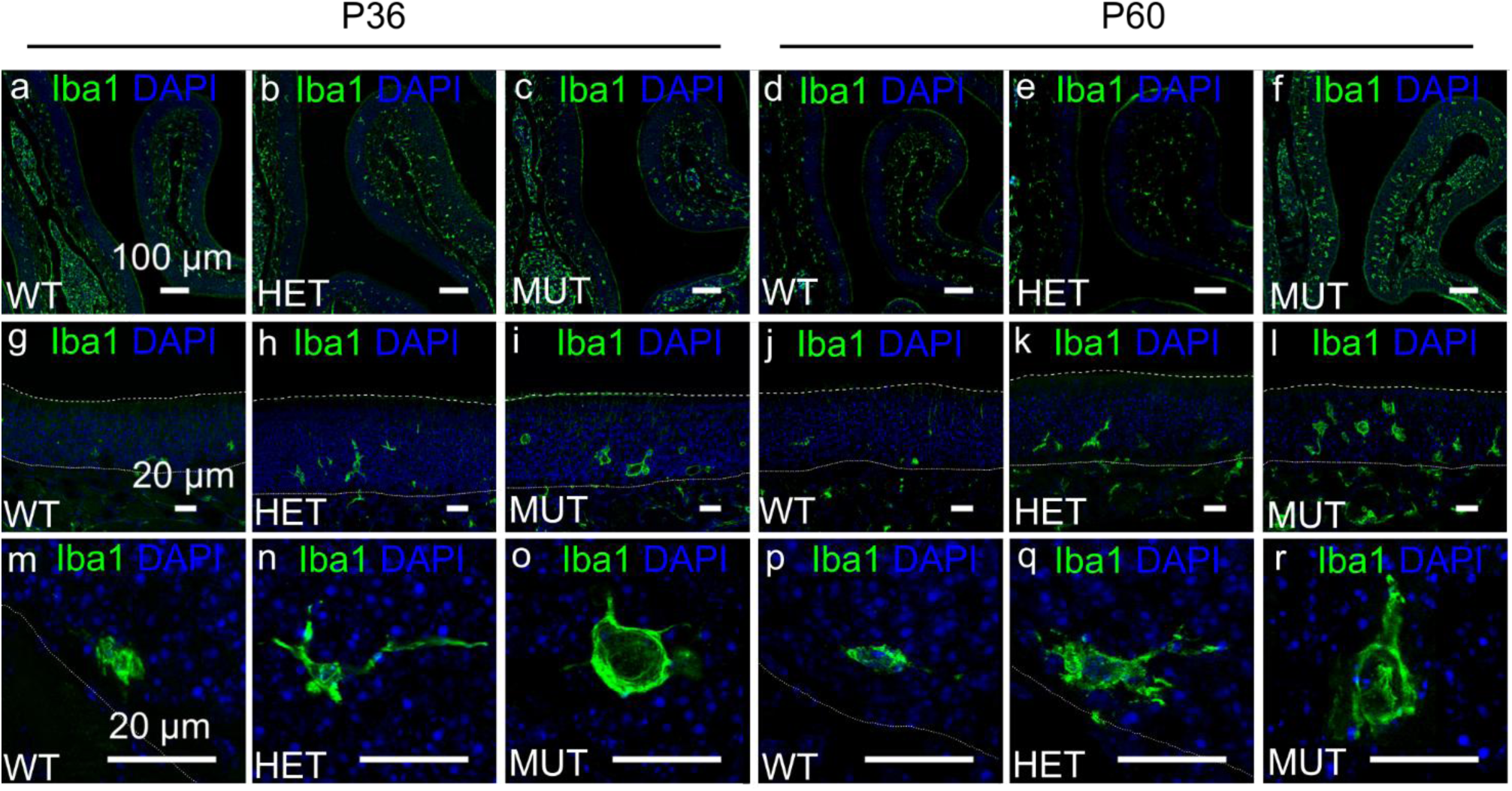
Pronounced microgliosis and macrophages recruitment in OE of MUT mice. Immunohistochemical reaction of Iba1+ microglial cells in the OE (a – f: g – l), (g – r) the white dotted line represents the OE basal layer while the white dashed line marks the ciliary layer. Coronal sections of HET (b) and MUT (c) mice revealed more reactivity and infiltration of Iba1+ cells compared to WT at P36 (a). MUT mice at P60 (f) demonstrated a noticeable increase of Iba1+ cells in in the OE compared to WT (d) and HET (e) mice. At P60 activated macrophages appear clustered more densely in OE (c, f) than below the basal lamina of MUT mice. At P36 (i, l) and at P60 (l, r) macrophages show a rounded shape in MUT mice compared to WT (g, j, m), HET mice (h, k, n, q) show an intermediary phenotype with ramified processes. Sample size: WT = 2 mice, HET = 1, MUT = 2 at P36; sample size: WT = 3 mice, HET = 1, MUT = 1 at P60.

### Altered phospholipid profiles in OE isolated from NPC1 MUT mice

Using Iba1 as a marker, we showed that neuroinflammatory processes are active in the OE of MUT mice as early as P36. We decided to look for possible lipid biomarkers of inflammation and performed comparative lipid analysis of OE isolated from WT, HET and MUT mice by positive ion mode MALDI-TOF/MS.

Figure 4 a illustrates the representative lipid profiles of WT and MUT OE at P60, where the main peaks in the mass spectra correspond to the [M+H]^+^ molecular ions of choline headgroup-containing glycerophospholipids. By comparing the two panels (upper and lower in ©. 4 a, both the lipid profiles, although not perfectly overlapping, share many similarities and no qualitative differences can be observed between the phospholipid patterns in OE of WT and MUT mice. In both the mass spectra major signals are detected in the higher *m/z* range, at *m/z* 734.5, 760.5, 782.6 and 810.5, attributable to the [M+H]^+^ molecular ions of four phosphatidylcholine (PC) species (PC 32:0, PC 34:1, 36:4, 38:4, respectively). Smaller MALDI signals at *m/z* 496.6 and 524.4 are visible in both the OE mass spectra and are assigned to the [M+H]^+^ molecular ions of lysophosphatidylcholine (LPC) species (Fig. 4a). At *m/z* 496.6, the signal is attributable to the lysophospholipid LPC with a palmitic acid (LPC 16:0), while that at *m/z* 524.4 is assigned to LPC containing a stearic acid (LPC 18:0). MALDI lipid profiles of OE isolated from HET mice at P60 as well as those obtained from WT, HET and MUT mice at P36 (not shown) have similar profiles as the WT and MUT at P60 (shown in Fig. 4 a).

**Figure 4.**
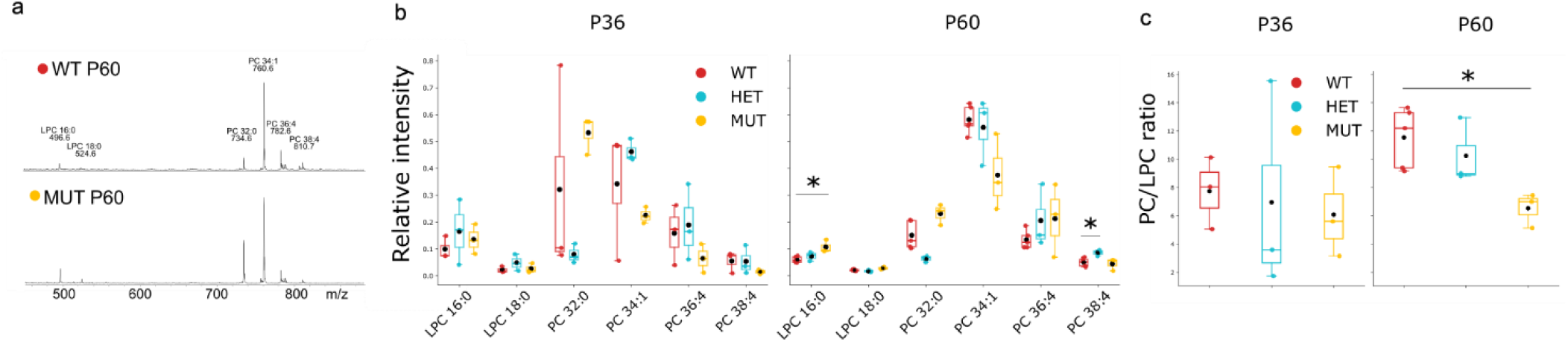
A lipid marker of neuroinflammation in OE of MUT mice. (a) Comparative lipid analysis of OE by MALDI-TOF/MS in WT (upper panel) and MUT mice (lower panel) at P60. Representative (+) mass spectra and lipid assignments for main peaks are shown. LPC = lysophosphatidylcholine; PC = phosphatidylcholine. (b) Differences in relative intensities of LPC and PC signals in OE mass spectra from WT, HET and MUT at P36 (left panel) and P60 (right panel). (c) PC/LPC ratio calculated in OE mass spectra from WT, HET and MUT at P36 (left panel) and P60 (right panel). The central black point represents the mean, central line: median, upper and lower box boundaries: 25th and 75th percentile, extreme lines: The highest and lowest value. Sample size: WT = 3 mice, HET = 3, MUT = 3 at P36; sample size: WT = 5 mice, HET = 3, MUT = 3 at P60.

Histograms in Figure 4 b show the statistical analysis of MALDI signal intensities obtained by comparing series of replicates of OE mass spectra from WT, HET and MUT, at P36 and P60, where alterations in acyl composition of phospholipids between groups of animals can be observed. No significant differences in signals assigned to different PC and LPC species were found, except for the peak diagnostic for LPC 16:0 that was significantly higher in the MUT lipid profile than in WT at P60 (Fig. 4 b). Interestingly, LPC levels are generally higher in many inflammatory states (Liu et al. 2020). Also, the signal assigned to PC 38:4 was significantly higher in HET, but not the MUT, lipid profile than in WT at P60 (Fig. 4 b).

Figure 4 c illustrates the PC/LPC ratios calculated from the various MALDI OE mass spectra of WT, HET and MUT at P36 and P60. This ratio in OE from the MUT mice at P60 was significantly lower than that calculated in WT. Notably, a lower PC/LPC ratio is generally a sign of increased inflammation (Fuchs et al. 2005; Angelini et al. 2014; Palazzo et al. 2025).

In conclusion comparative lipid analyses of OE obtained from different groups of animals revealed a significant enrichment in LPC species containing palmitic acid and markedly lower PC/LPC marker in OE from MUT mice at P60 compared with WT.

### OE stem cell niche

The horizontal basal cells (HBCs) are the OE multipotent reserve stem cells. They are flat cells which directly contact the basal lamina and define the boundary between the olfactory epithelium and the underlying lamina propria In physiological conditions, HBCs are mitotically quiescent, while in pathological conditions, they become more active and can regenerate the different cell types of the OE (Huard and Schwob 1995; Leung et al. 2007a; Iwai et al. 2008; Gadye et al. 2017). We used antibodies against cytokeratin 14 (CK14) as an HBC marker and found that only at P60 a difference in HBC density emerged in MUT (Fig. 5 f) compared to WT mice. At P60 also HET mice (Fig. 5 e, summary in g) showed a significant increase in HBC density compared to WT (Iwai et al. 2008; Schwob et al. 2017; Child et al. 2018) (Fig. 5 c, f).

**Figure 5.**
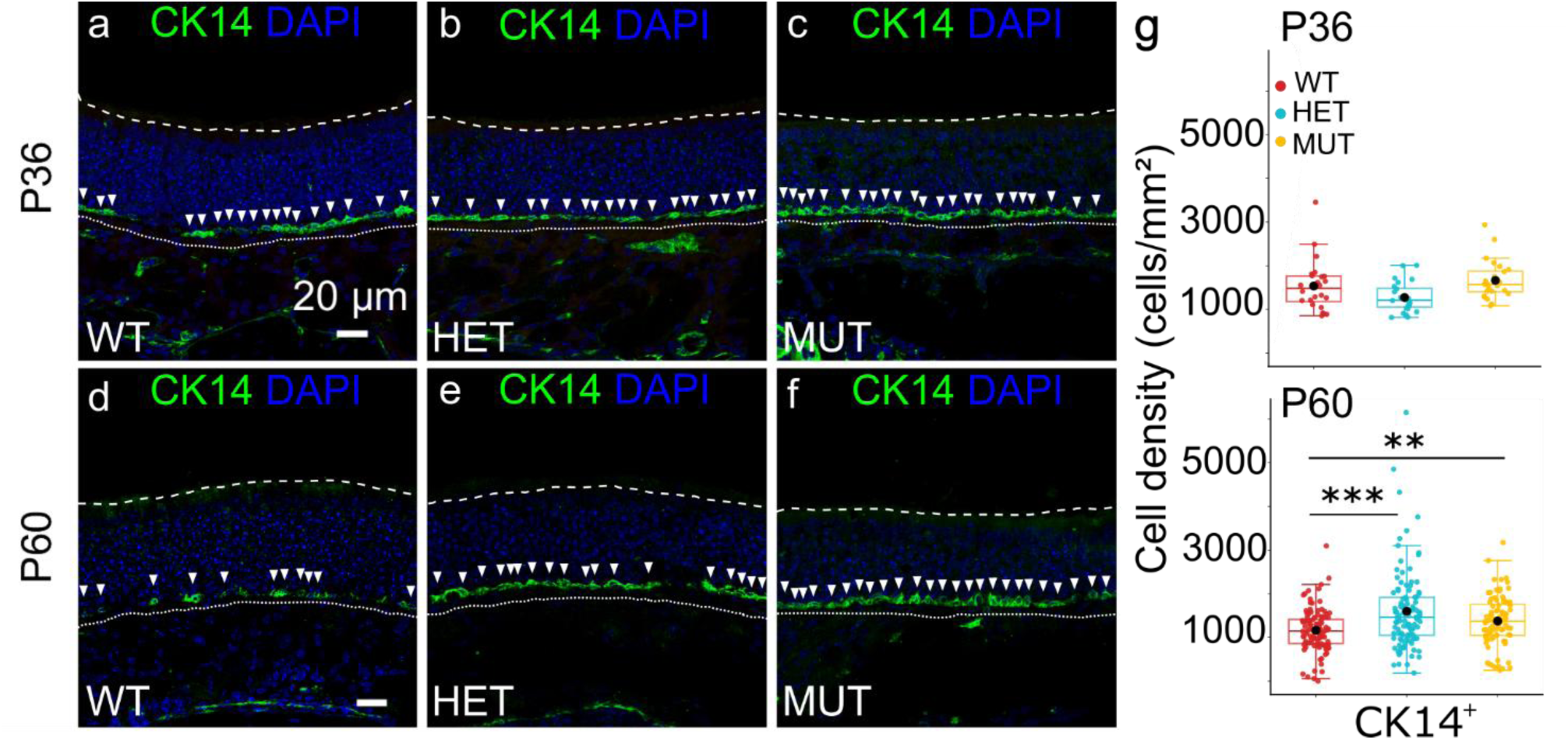
HBCs density increases in MUT mice at P36 and P60. (a – f) Coronal sections from WT, HET and MUT olfactory epithelium showing HBCs (CK14+) in mice ages P36 and P60. HBCs were counted considering the DAPI nuclei surrounded by the CK14 green staining as indicated by the white arrowheads. They lie close to the basal lamina (white dotted line). (g) Box plots showing cell density in WT, HET and MUT mice. The central black point represents the mean, the central line: the median, upper and lower box boundaries: 25th and 75th percentile, extreme lines: the highest and lowest value. Kruskal–Wallis test followed by Mann-Whitney and Benjamini–Hochberg (BH) post hoc analysis was performed to CK14+: H(2) = 10.32, p = 5.73e-3 at p36 and at p60: H(2) = 19.06, p = 7.26e-5; p = **0.01, ***0.001. Sample size: n = 3 mice for each genotype and each age. For more detailed on the post-hoc analysis refer to the table 1.

The globose basal cells (GBCs) are a heterogeneous population that can be identifyed by the different transcription factors that they express (Schwob et al. 2017). We evaluated the proliferative subpopulation of GBCs by using antibodies against Ki67, a transcriptional factor used as a marker of active cell cycling (Schnittke et al. 2015) in S phase. Most of the cells labeled by anti-Ki67 (Fig. 6 white arrows) are presumably proliferative GBCs and additional Ki67^+^ are duct/gland cells and sustentacular cells (Fig. 6 yellow arrowsheads) (Jang et al. 2014). Ki67^+^ basal cell (GBCs) density does not change in MUT and HET mice compared to WT at P36 (Fig. 6 a, b, c) and P60 MUT (Fig. 6 d, e, f).

**Figure 6.**
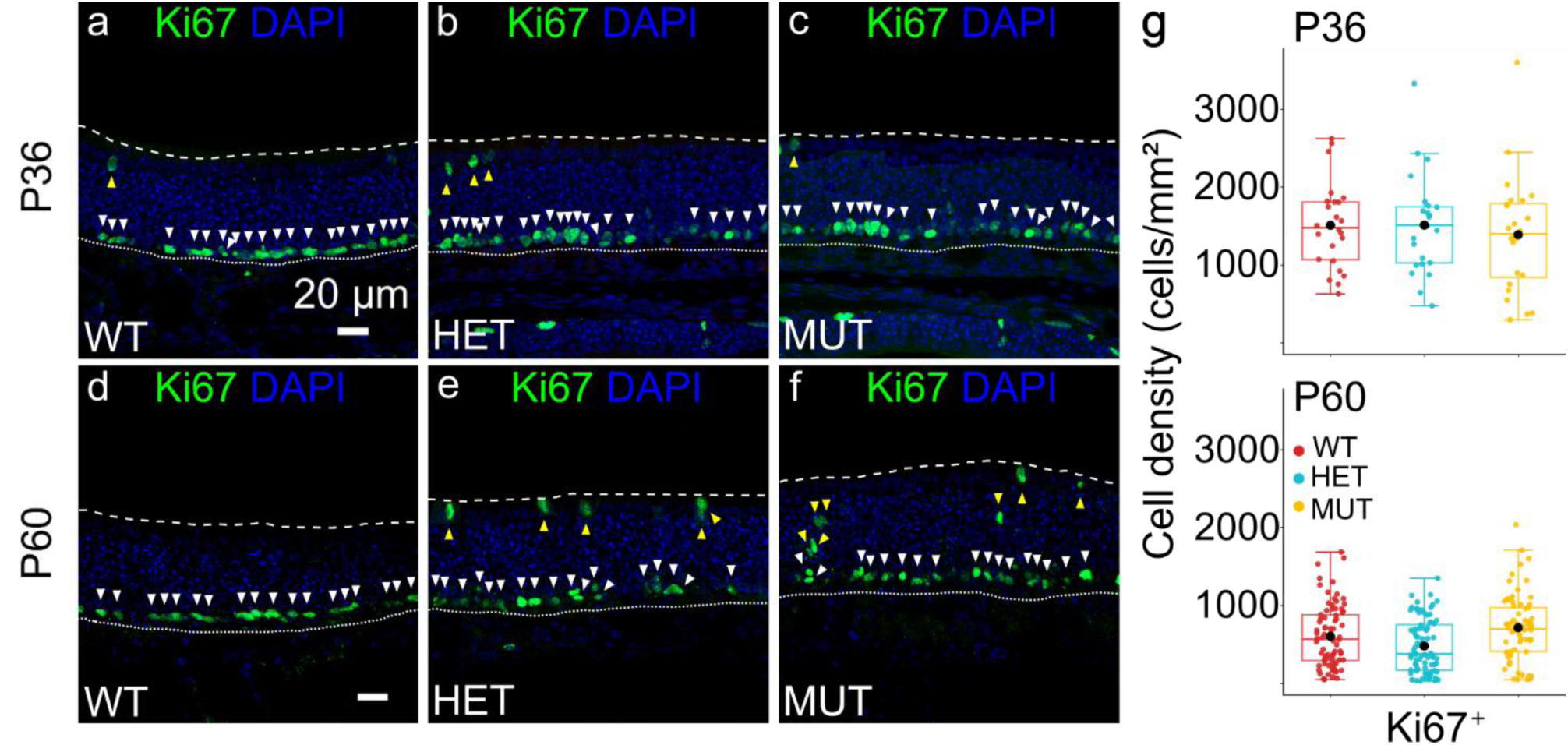
Basal proliferating staminal cells density increases in MUT mice at P60. (a – f) Coronal sections from WT, HET and MUT olfactory epithelium in mice ages P36 and P60. Proliferating stem cells were counted considering the basal (basal lamina indicated by white dotted line) Ki67+ cells and they represent proliferative GBCs and/or HBCs (white arrowheads). Yellow arrowheads showing non basal proliferating cells closer to the ciliary layer (white dashed line). (b) Box plots showing cell density in WT, HET and MUT mice. The central black point represents the mean, the central line represents the median, upper and lower box boundaries: 25th and 75th percentile, extreme lines: The highest and lowest value. Sample size: n = 3 for each genotype and for each age. For more detailed on the post-hoc analysis refer to the table 1.

### EOG responses are reduced in MUT mice

We asked whether the structural differences that we found were causing functional alteration of the OE. Therefore, we performed air-phase electro-olfactogram (EOG) recordings from the OE of WT, HET and MUT mice at both P36 and P60. When stimulated with isoamyl acetate (IAA) at 10^-1^ M, we observed a response that was, on average, around -11 mV in WT and that decreased to about half in MUT mice at P36 (Fig. 7 a) and P60 (Fig. 7 c). We stimulated the OE using different concentrations of IAA (10^-3^ M, 10^-5^ M, 10^-7^ M) and we could again observe a decrease in response amplitude in MUT animals across concentrations. Dose-response plots showed that the amplitude of the response was overall smaller in the MUT mouse models compared to the WT at both P36 (Fig. 7 b) and at P60 (Fig. 7 d). The reduction in odorant responses may be due to the decreased density of OMP^+^ neurons in the OE of MUT mice.

**Figure 7.**
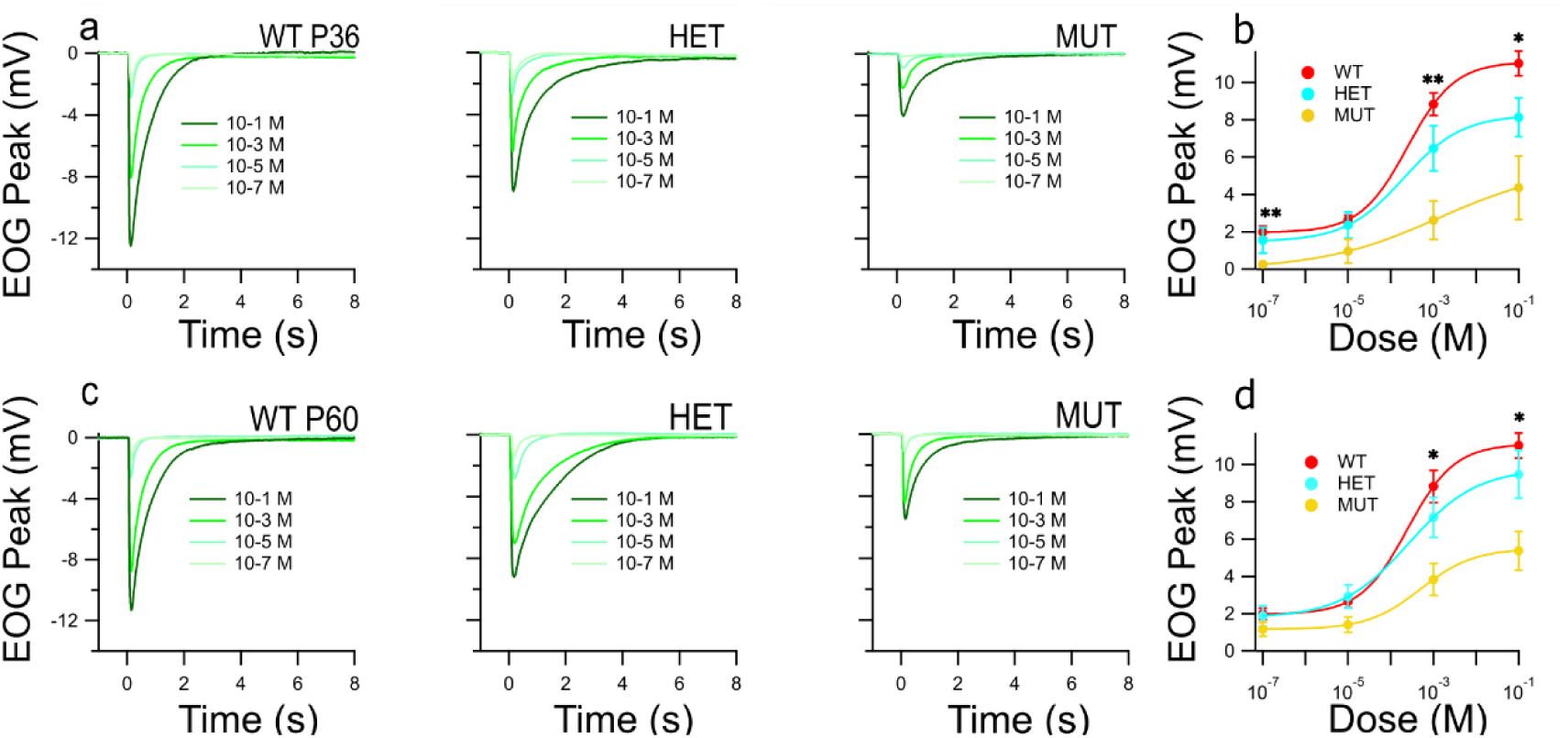
OSN neurodegeneration affects odorant-evoked responses. Odorant-evoked EOG responses to 100 ms stimulations with isoamyl acetate (10-1 M, 10-3 M, 10-5 M, 10-7 M) were recorded from turbinate IIa of WT, HET and MUT mice at P36 (a) and P60 (c). Summary of EOG dose-responses isoamyl acetate in MUT HET and WT mice at P36 (b) and at P60 (d). Data are presented as mean and SEM. A linear mixed model performed on EOG responses to different concentrations revealed genotype, dose, and genotype-dose interaction as significant effects (genotype: F(2,17) = 8.43, p = 2.86e-3, dose: F(3,51) = 172.53, p = 4.32e-10, genotype-dose: F(6,51) = 8.78, p = 1.41e-6 at P36; genotype: F(2,14) = 4.25, p = 3.61e-2, dose: F(3,42) = 209.3, p = 3.09e-9, genotype-dose: F(6,42) = 9.44, p = 1.46e-6 at P60; p = *0.05, **0.01. Sample size: WT = 9 mice, HET = 8, MUT = 5 at P36; WT = 7 mice, HET = 6, MUT = 4 at P60. For more detailed on the post-hoc analysis refer to the table 1.

### Olfactory-driven behaviors are impaired in MUT mice

We next asked whether our findings are behaviorally relevant. We sought to investigate how the observed changes combine to alter naïve behavior such as tracking odors to find hidden food. We performed an odor-guided food-seeking test consisting of a hidden piece of cookie under the cage bedding (Machado et al. 2018; Meyer et al. 2018; Mitrano et al. 2021; Lobraico et al. 2025). The mouse is free to explore the cage, and the latency to find the cookie is measured. MUT mice took more time to find the cookie than WT and HET at P36 (Fig. 8 left panel). MUT mice took more time to find the cookie also at P60 (Fig. 8 right panel) compared to WT. We did not find differences across the genotypes when the cookie was placed on top of the bedding, when the animals could easily reach it when they did not have to rely on olfaction to find the cookie, excluding gross motor disorders or alterations in the motivation to find the food.

**Figure 8.**
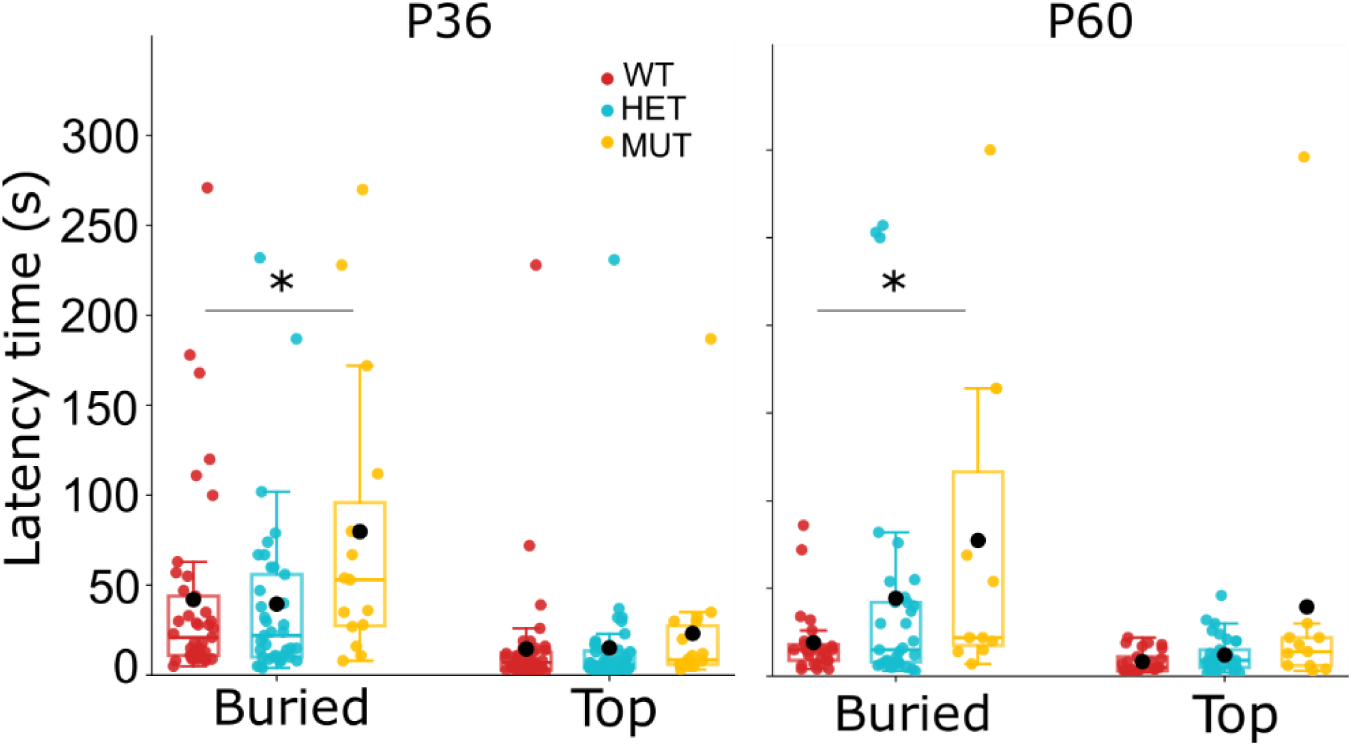
OSNs neurodegeneration affects sense of smell. Latency time in finding the buried cookie or placed on top of the bedding at P36 (left panel) and at P60 (right panel). MUT mice are slower than WT in finding the cookie. The black central point in the box plots represents the mean, central line: median, upper and lower box boundaries: 25th and 75th percentile, extreme lines: the highest and lowest value. Kruskal–Wallis test followed by Benjamini-Hochberg (BH) post hoc analysis was performed on buried cookie: H(2) = 5.63, p = 5.99e−2 at P36; H(2) = 5.88, p = 5.28e−2 at P60; p = *0.05. Sample size: WT = 41 mice, HET = 37, MUT = 15 at P36; sample size: WT = 25 mice, HET = 34, MUT = 11 at P60. Mice that did not find the cookie were excluded from the analysis. For more detailed on the post-hoc analysis refer to the table 1.

### Altered sense of smell in NPC1 patients and their heterozygous parents

Do patients with NPC1 show olfactory impairments? To the best of our knowledge, the olfactory abilities of the NPC1 patients have not been reported yet. We had the opportunity to perform olfactory testing on a family where the child was carrying two mutations of the NPC1 gene that were classified as pathogenetic. Each of the parents was heterozygous for one of the mutations. The child and the parents did not report any clear sensory deficits.

We used the Sniffin’ Sticks test (Hummel et al. 1997; 2007) to test the olfactory abilities of the family and to compare our results with the normative data published by Oleszkiewicz et al (Oleszkiewicz et al. 2019). The child (age 21) had a TDI (Threshold, Discrimination, Identification) score of 19.5, which is much below the 5^th^ percentile of their age range (age 21-30, 5th perc. = 29.96) or below the cut-off for hyposmia of 30.75 (Oleszkiewicz et al. 2019). All three subtests of the Sniffing’ Tests (Threshold (T) = 5.5, Discrimination (D) = 5, Identification (I) = 9) were below the 10^th^ percentile of their age group, thus we could classify the patient as hyposmic. Interestingly, the mother (heterozygous for one of the mutations in the <u>Npc1</u> gene, age 50) scored 26.25, which is below the cut-off for hyposmia. Her subtest results were T = 5.25, D = 11, I = 10. The father scored 30.5 (T = 5.5, D = 13, I = 12, age 51), which although close, was also below the 30.75 cut-off for hyposmia.

## 2. Discussion

NPC1 is a rare autosomal recessive lipid storage disease that causes progressive neurodegeneration. We decided to use the B6.129- *Npc1^tm(I1061T)Dso^* mouse model with the I1061T missense mutation in the *Npc1* gene as it represents the most frequent mutation observed in human patients (Carstea et al. 1997; Millat et al. 1999; Gelsthorpe et al. 2008). The murine NPC1I1061T protein has a reduced half-life in vivo, consistent with protein misfolding and rapid ER-associated degradation. Here, our results showed that neurodegeneration in OE is impairing olfactory functions as early as P36, making it potentially a sensitive marker for NPC1 disease progression.

### Early OSNs neurodegeneration in NPC1 disease

We found that in our *Npc1^tm(I1061T)Dso^ homozygous mouse* model neuronal degeneration rather than a failure to generate immature neurons caused a significant reduction of OMP^+^ mature OSN density at P36 in MUT mice relative to WT and HET, while the density of Tuj1^+^ immature OSNs is preserved. This extends prior NPC1 work showing structural and functional deficit in the olfactory system in a knock out mouse model (Hovakimyan et al. 2013; Seo et al. 2014; Meyer et al. 2018; Dragotto et al. 2019; Rava et al. 2022). We showed that NPC1 protein stability is important to maintain cell integrity in the OE. Cholesterol-trafficking defects can selectively impair mature OSNs while leaving almost intact the regenerative abilities of the OE. The persistence of immature neurons suggests that although neurogenesis remains active, newly generated OSNs fail to mature or survive long-term. This trajectory mirrors the progressive cell loss described in other neural regions in NPC models, particularly Purkinje cells in the cerebellum (Praggastis et al. 2015a) and other sensory systems affected by NPC, including the retina (Palladino et al. 2015) and auditory pathways (Ko et al. 2005; Pressey et al. 2012). Our results suggest that the ongoing apoptosis in the OE stimulate the HBCs to differentiate to compensate for neuronal loss. The persistent neuroinflammatory processes in the OE may exacerbated the entire loop. Crucially, these results match other neurodegenerative diseases (Klein et al. 2021; Son et al. 2021; Axmann and Lehrner 2025) like Alzheimer’s disease (Huart et al. 2015; Attems et al. 2021) and Parkinson’s disease (Tremblay et al. 2024; Chowdhury et al. 2025), which also show early olfactory decline. This further supports the notion that mature *Npc1* MUT OSNs are vulnerable targets across neurodegenerative diseases no matter the etiology. OMP-based readouts may be applicable in humans and would provide a sensitive and accessible tool to investigate early neurodegenerative change in the OE.

The proliferative GBCs does not change in MUT mice compared to WT indicating that the OE stem cell niche is different from other neurogenic systems, such as the hippocampal dentate gyrus or the subventricular zone, where neurodegeneration often results in progenitor depletion and impaired regenerative capacity (Winner et al. 2011; Moreno-Jiménez et al. 2019). The maintenance of GBC function in MUT mice indicates that the OE stem cell niche is remarkably resilient to cell-autonomous damage related to cholesterol trafficking impairments even during advanced disease stages.

HBCs are immunoreactive for cytokeratin 14 (CK14^+^) (Getchell et al. 2000) and by P60 we could observe an increase in their density in HET and MUT mice. HBCs in MUT mice acquire a spherical morphology which is more evident and consistently present than in WT mice. This morphological rearrangement could reflect a transition from quiescence to a primed or activated state (Schwob et al. 1995; Leung et al. 2007b; Durante et al. 2020; Gameiro et al. 2025), consistent with previous evidence showing that HBCs undergo structural remodeling prior to engaging in regeneration (Gadye et al. 2017). The persistence and even expansion of the HBC pool in *Npc1* mutants may thus represent a compensatory response to ongoing degeneration, preserving the regenerative potential of the OE since HBCs can give rise to GBCs too and eventually to new OSNs (Iwai et al. 2008). In previous studies using *Npc1* mouse models, the OE showed increased proliferation (GAP43 / BrdU) despite severe OSN loss indicating that the GBC/HBC niche can be active (Meyer et al. 2017; Dragotto et al. 2019). The decreased density of mature OSNs together with the altered odorant responses measured in the EOG, indicate that in MUT mice the rate of apoptosis is faster than that of the turnover. Our data demonstrate that apoptosis in the OE increases across disease stages in the *Npc1* mouse, with a clear rise in cCasp3⁺ OSNs at P36 and P60 in MUT and, by P60, a detectable elevation was also observed in HET. Our results on cCasp3 confirms the cell death pathway by effector caspases is a recurring theme of neurodegeneration-linked olfactory decline as reported for Npc1 disease or eosinophilic chronic rhinosinusitis (CRS) (Hetmańczyk-Sawicka et al. 2020; Chen et al. 2024).

### Npc1 misfolded protein causes increased neuroinflammation and microglia activation during disease progression

It has been suggested that neurodegeneration in NPC1 disease begins in a neural cell-autonomous and microglia-independent manner (Loftus et al. 1997; Peake et al. 2011). Therefore, the apoptotic events are triggered by the impaired egress of cholesterol and sphingolipids from late endosomes/lysosomes in neurons can lead to mitochondrial stress, (Liedtke et al. 2022), each of which can drive caspase activation (Cai et al. 1998; Wu et al. 2005; Huang et al. 2006; Wheeler and Sillence 2020). Nevertheless, in the OE of the *Npc1* mouse model used here microglia activation / macrophages recruitment and infiltration is more visible in MUT and HET mice compared to WT. Iba+ cells infiltrating the OE may be the results of OSN cell death and in this context inflammation might exacerbate NPC1 disease (Cougnoux et al. 2023). Microgliosis is a common finding in the olfactory bulb (OB) of olfactory-deficient animals, including NPC1 mouse models (Hovakimyan et al. 2013; Seo et al. 2016).

### Olfactory functions are impaired in MUT mice as early as one month of age

The structural changes we observed in the OE explain the functional impairments we observed in EOG experiments. We performed dose response EOG recordings and recorded a reduction in response amplitudes at different odorant concentrations rather than change in sensitivity in line with the idea that it is determined by the reduced mature OSNs density in MUT mice. Previous reports (Hovakimyan et al. 2013; Meyer et al. 2018) showed deficit in EOG recordings in a knock out mouse model for Npc1 but lacked both the dose responses and investigation during disease progression by focusing on a single time windows.

In our behavioral assay, MUT took longer to find the cookie compared to WT. The contribution of the peripheral olfactory system in such behavior is crucial since altered transduction in OSNs negatively alters the ability of mice to perform the test (Klein et al. 1996; Pietra et al. 2016; Rava et al. 2022). Since the test does not rely on an operant-conditioning task, mice must use their innate sense of smell to locate the odor source food. Previous papers that studied olfaction in *Npc1* knock out mouse models showed altered olfactory behavior (Meyer et al. 2018; Dragotto et al. 2019; Lyons-Warren et al. 2021; 2023). Regarding our phospholipid profiling of OE from NPC1 mice, it is well known that LPC levels can increase in many inflammatory diseases, and a PC/LPC ratio decrease is often used as an indicator of membrane integrity disruption and neuronal damage (Mulder et al. 2003; Fuchs et al. 2005; Angelini et al. 2014). LPC acts as a pro-inflammatory mediator, attracting immune cells like neutrophils and T cells, and by stimulating the release of inflammatory factors, while high PC levels are generally associated with anti-inflammatory effects (Balood et al. 2014; Zahednasab et al. 2014; Liu et al. 2020). Here lipid analysis revealed a significant enrichment in a LPC species and markedly lower PC/LPC ratio in OE from MUT mice at P60 compared to WT and therefore we propose to extend this test to NPC1 patients and to their parents by performing MALDI-TOF lipid analysis on biopsies taken from their noses.

### Smell test as part of the tools for neurological evaluations in NPC1 disease

We described the pathophysiological and functional changes occurring in the olfactory system in a mouse model closely resembling NPC1 in humans (Millat et al. 1999; Vanier and Millat 2003; Vanier 2010; Hovakimyan et al. 2013; Meyer et al. 2018). We then sought to investigate whether olfactory impairments are present in patients. We performed olfactory testing with a family of three with the parents carrying two different heterozygous mutations and their child carrying both parental mutations. The child showed a TDI score that pointed at severe hyposmia (Table 2). Similar results were found in at least one of the parents although less severe than that of the child. Interestingly, all the three subjects showed low scores in the Threshold test, which should point more at peripheral damage rather than central along the olfactory pathways. These observations may suggest that peripheral damages are the first to occur not only in homozygous but also in the heterozygous subjects. Therefore, if the periphery is altered at early age, we may look for biomarkers in i.e. nasal biopsies that are easily accessible.

**Table 1.**
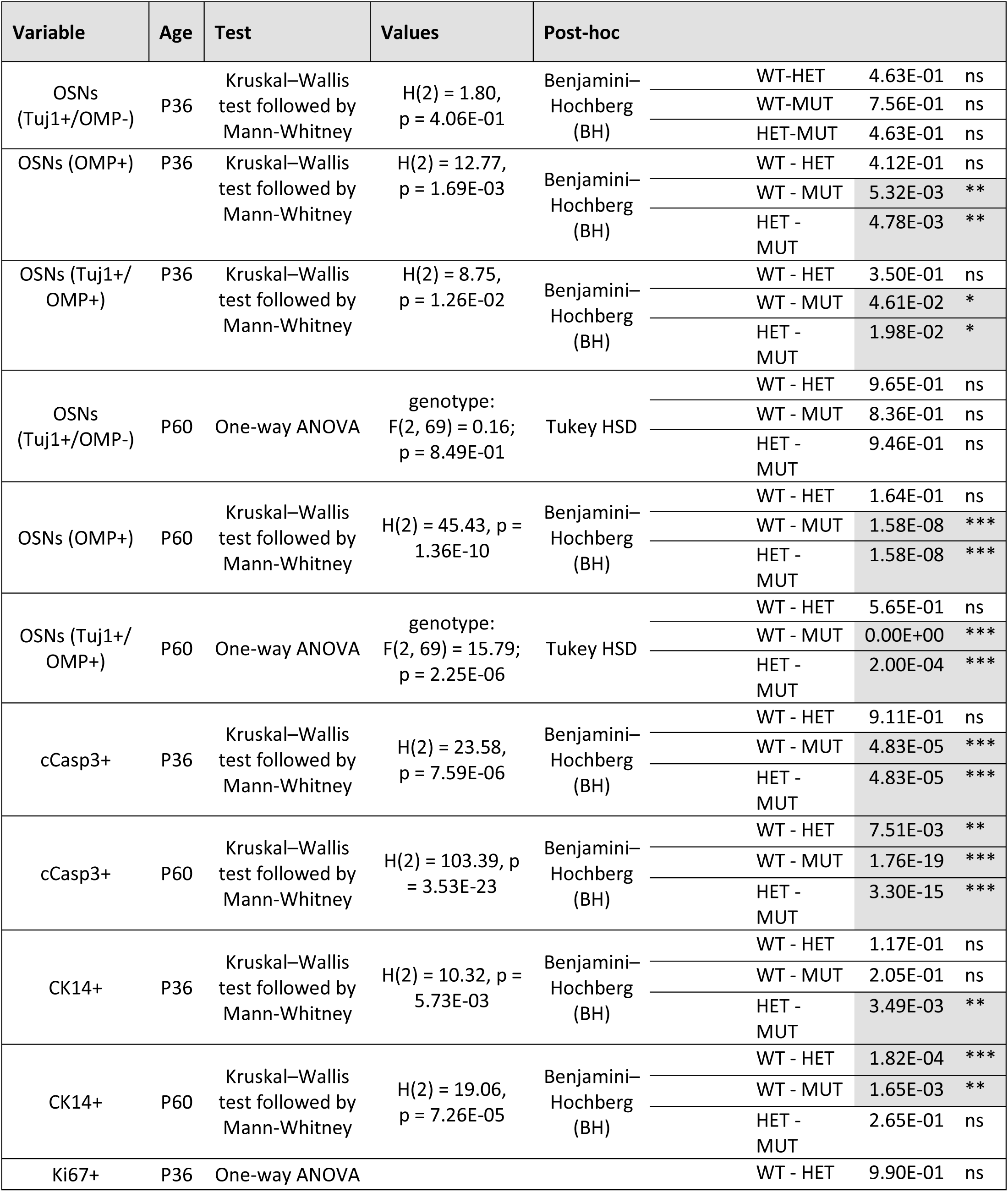
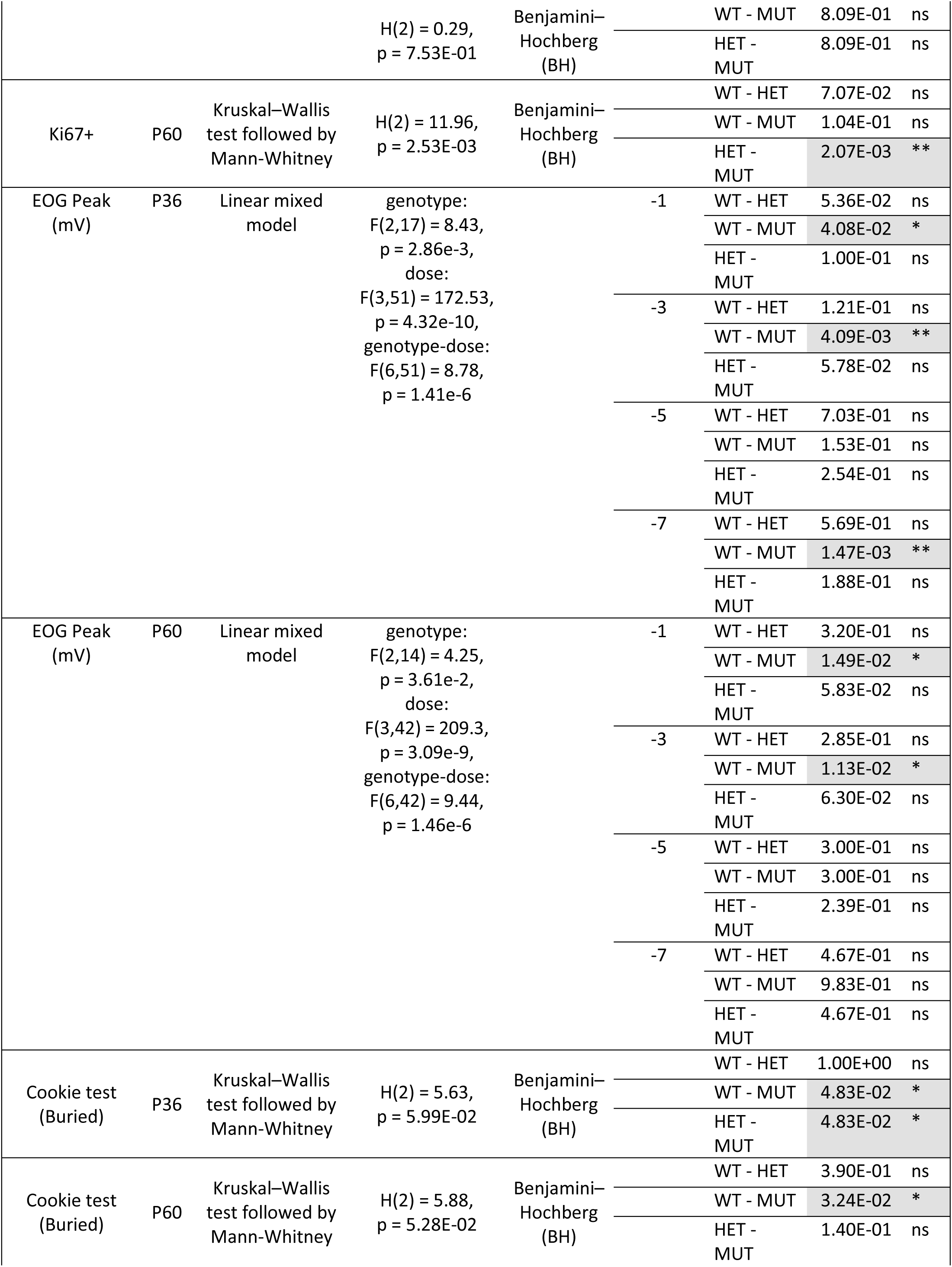
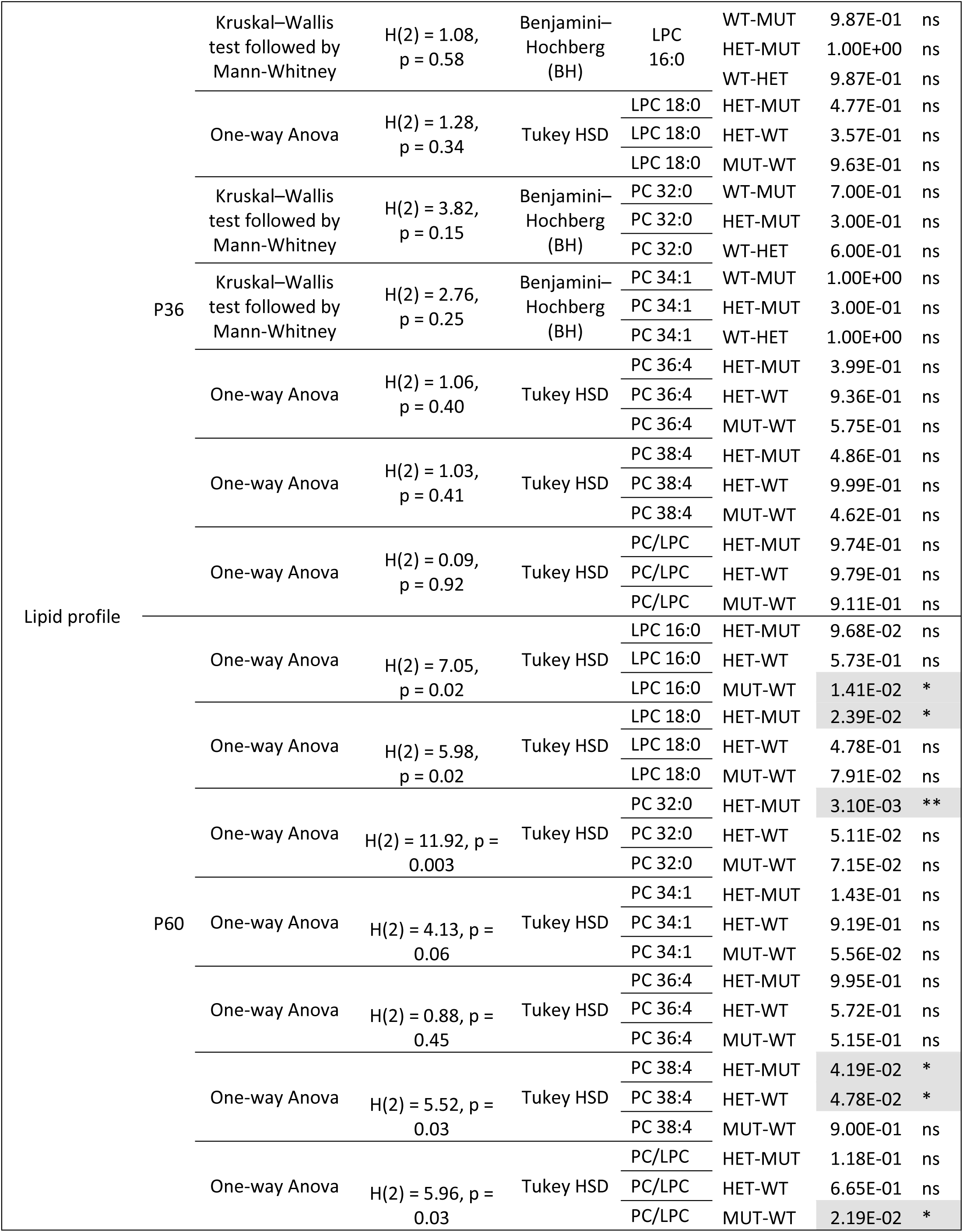
Summary statistical analysis

**Table 2.** Sniffin’ sticks score of the NPC1 child and their parents. Cut-off values for hyposmia is 30.75 according to Oleszkiewicz et al 2019.

|  | Threshold | Discrimination | Identification | TDI | Notes |
| --- | --- | --- | --- | --- | --- |
| NPC1 patient | 5.5 | 5 | 9 | 19.5 | Severe Hyposmia |
| Mother (Het) | 5.25 | 11 | 10 | 26.25 | Hyposmia |
| Father (Het) | 5.5 | 13 | 12 | 30.5 | Mild Hyposmia |

## Conclusions

In our *Npc1* mouse model, abnormalities in intracellular cholesterol transport with subsequent accumulation of lipids in the OE causes OSN neurodegeneration, microglia activation and increased apoptosis very early during the disease progression. These alterations at a cellular level have deep consequences on olfactory abilities as demonstrated by electrophysiological recordings and behavioral tests, the latter phenotype being similar to those observed in human NPC1 disease. Thus, the loss of the sense of smell could be used as an early and easily accessible sentinel symptom to diagnose the disease and guide treatments and approaches that prevent/slow neurodegeneration while simultaneously ameliorate olfactory dysfunction.

Finally, we showed that NPC1 homozygotic patients may be affected by olfactory impairment even in the absence of any other clear sensory and neurological signs of the disease. In addition, heterozygosity may also alter sense of smell. A limitation of our study is given by the low number of our subjects, but it is worth considering that NPC1 is a rare disease, and it is not easy to recruit young or young adults that could complete the tests. Nonetheless, our results from parents should suggest that olfactory testing should be included in the panel of the neurological examinations since heterozygosity itself has been implicated in fostering neurodegeneration, adding another layer of complexity to the disease’s understanding.

## Materials and Methods

### Animals

The mouse model B6.129- *Npc1^tm(I1061T)Dso^* with the I1061T missense mutation in the *Npc1* gene were generated by the Jackson Laboratory JAX stock #027704 (Bar Harbor, ME USA) as previously described by (Praggastis et al. 2015). Mice were maintained under a light/dark cycle (12/12 h) in the Department of Translational Biomedicine and Neuroscience’s animal facility at room temperature (22±2°C) and 75% humidity, with food and water ad libitum. All experiments were performed using procedures approved by the Institutional Committee on Animal Research and Ethics of the University of Bari and the Italian Health Department (Project n°465/2022-PR) and in accordance with the European directive on animal use for research. Experiments were carried out on mice from 1 to 2 months old and of either sex. All experiments were designed to minimize the number of animals used and their suffering.

In order to genotype B6.129- *Npc1^tm(I1061T)Dso^*, a 0.5 mm tail was collected, and DNA was extracted using the PCR kit (Thermo Scientific Phire Animal Tissue Direct PCR Kit, Thermo Fisher Scientific, USA). The extracted DNA was amplified by PCR using specific primers: Mouse Forward 24238: 5′- TCA GCC CTC TCT TAC TTG GTG -3′ Mouse Reverse 24239: 5′- CTG CTG TGT GCT AGG AAT CAC T -3′ The 210 bp PCR product represents the allele with the mutation while the 146 bp is the wild type (WT) allele. The amplified DNA produce one 210 bp band in case of homozygosity for the mutation, two bands (210, 146) in case of heterozygosity and one band of 146 bp in case of homozygosity for the wild type allele.

### Immunohistochemistry

Mice were sacrificed by cervical dislocation after general anesthesia with CO2. The dissected nasal tissues were fixed overnight at 4°C in 4% PFA and then rinsed in PBS Ca²⁺/Mg²⁺ (pH 7.4). Decalcification was performed in 0.5 M EDTA (pH 8) over several days to a week at 4°C, with periodic replacement of the buffer; the duration depended on the bone mineralization. Following PBS washes, tissues were sequentially immersed in sucrose solutions of 5%, 10%, and 20% (w/v) in PBS Ca²⁺/Mg²⁺ for 10 minutes each, and then incubated overnight in 30% sucrose at 4°C. After rinsing, samples were embedded in Tissue-Tek OCT compound (Sakura, The Netherlands) and frozen. Coronal sections, 14–15 μm thick, were cut using a cryostat (CM 1900, Leica) at −20°C and stored on poly-L-lysine-coated slides at −80°C.

For immunostaining, sections were rehydrated in PBS Ca²⁺/Mg²⁺ and subjected to antigen retrieval with 0.5% SDS (w/v) in PBS Ca²⁺/Mg²⁺. Blocking was performed for 30 min to 1 h using either 5% normal goat serum, 4% BSA, 5% non-fat dry milk, 0.3% Triton X-100; or 5% normal donkey serum with 4% BSA in 0.3% Triton X-100; or 5% BSA with 0.3% Triton X-100, all prepared in PBS Ca²⁺/Mg²⁺. Sections were then incubated overnight at 4°C with primary antibodies diluted in the blocking solution, including: goat anti-OMP (1:300; FUJIFILM Wako Chemicals U.S.A. Corporation, Richmond, USA), mouse anti-Tuj1 (1:300; BioLegend, San Diego, California.), rabbit anti-Cytokeratin14 (1:300; Proteintech, Rosemont, USA), rabbit anti-Ki67 (1:300; Abcam, Cambridge, UK), rabbit anti-cleaved Caspase-3 (1:100; Cell Signaling Technology, Danvers, USA), and rabbit anti-Iba1 (1:200; FUJIFILM Wako Chemicals U.S.A. Corporation, Richmond, USA).

The following day, secondary antibodies AlexaFluor 488 goat anti-rabbit, AlexaFluor 594 donkey anti-goat, AlexaFluor 488 rabbit anti-mouse, or AlexaFluor 555 donkey anti-rabbit (all from Life Technologies, Thermo Fisher Scientific, Carlsbad, California, USA) were applied at 1:1000 in 0.2% Tween-20 in PBS Ca²⁺/Mg²⁺ for 45–60 min at room temperature. After washing, sections were mounted with either a 1:1 mixture of Mowiol-DAPI or VECTASHIELD Vibrance Antifade Mounting Medium. Imaging was performed with a Leica TCS SP8 confocal microscope using 20×/0.40 dry, 40×/1.30 or 63x/1.40 oil objectives at 1024×1024 pixels across the z-stack with zoom factor 1. Lower-magnification images were captured using a semi-automated fluorescence Leica stereomicroscope (M205FCA), using 2×/12.5 objective and all images were analyzed with Fiji software (Schindelin et al. 2012).

### Cell counting

The number of cells positive to OMP, Tuj1, CK14, Ki67, cCasp3 were counted from 8 to 12 random areas of coronal sections 14-15 μm thick, spanning widely separated regions in the olfactory epithelium. Cell density was measured in defined μm^2^ areas, that included all the OE thickness. Images were acquired with a LED fluorescence microscope (DM2500, Leica) using 40X or 20X/0.55 HC PL FLUOTAR objective at 1280 x 1024p and a CCD camera (DFC7000-T, LEICA), then analyzed with Fiji software.

### OE isolation

#### Matrix-assisted laser desorption ionization-time-of-flight/mass spectrometry (MALDI-TOF/MS) lipid analysis

MALDI-TOF mass spectra of OE samples were acquired on a Bruker Microflex LRF mass spectrometer (Bruker Daltonics, Bremen, Germany). All mass spectra were acquired in positive ion mode in reflector mode (detection range: 400–2000 mass/charge, *m/z*) using the delayed pulsed extraction. For each mass spectrum, 3000 single laser shots were averaged. External calibration with standard lipids (Avanti Polar Lipids from Alabaster, AL, United States) was performed, as previously described (Lobasso et al., J cell Physiol 2017; Lobasso et al., Front Physiol 2021).

OE lipids were analysed using a rapid lipid extraction protocol (mini-extract) suitable for small amounts of biological samples (Angelini R. et al. 2015). The matrix 9-aminoacridine hemihydrate (9-AA) was purchased from Acros Organics (Morris Plains, NJ, United States) and used at 10 mg/ml in 2-propanol/acetonitrile (60/40, v/v) for lipid analysis. 0.5 μL of lipid “mini-extract” obtained from OE (about 20 μg protein) was spotted on the MALDI target (Micro Scout Plate, MSP 96 ground steel target). The laser fluence was about 5% above threshold to have a good signal-to-noise (S/N) ratio. Peak areas, baseline correction, spectral mass resolutions and S/N ratios were determined by Flex Analysis 3.3 software (Bruker Daltonics, Bremen, Germany). Lipid Maps Database (http://www.lipidmaps.org/tools) was used for lipid identification. Electro-olfactogram recording The experimental setup followed procedures similar to those described previously by Zhao et al. (1998), Cygnar et al. (2010), and Lobraico et al. (2025). Mice were anesthetized using CO₂ and then decapitated. The heads were bisected sagittally along the midline and placed under a dissecting microscope (Stemi DV4, Zeiss) on a vibration-isolated table (Supertech Instruments), protected from electrical interference using a Faraday cage. The septum was removed to expose the turbinates, and the tissue was continuously perfused with distilled water bubbled at 37°C to maintain the hydration of the OE.

Recording electrodes were made from borosilicate glass (World Precision Instruments) using a P-1000 puller (Sutter Instrument) and positioned onto the surface of one of the turbinates. The pipette was filled with Ringer’s solution (140 mM NaCl, 5 mM KCl, 2 mM CaCl₂, 1 mM MgCl₂, 10 mM Hepes, 0.01 mM EDTA, pH 7.4 with NaOH) and the tip filled with 0.5% agar in Ringer’s solution. Odorant stimulation was delivered via a Picospritzer-controlled valve (Picospritzer II, Parker Hannifin) as a 100 ms vapor pulse at 10 psi into a continuous stream of humidified air maintained at 3 L/min (flowmeter, Masterflex). The Picospritzer was connected to the delivery tube, which carried odorant vapor generated from a 5 ml glass tube sealed with a rubber stopper and equipped with two 20-gauge needles serving as input and output ports. Isoamyl acetate (IAA) was prepared fresh daily, starting from a 5 M stock in DMSO (Honeywell, Riedel-de Haën) and diluted to final concentrations ranging from 10⁻⁷ to 10⁻¹ M in distilled water. Air/mucus partition coefficients were taken from Scott et al. (2014) and Coppola et al. (2017), with IAA having a value of −2.4038.

The reference electrode was placed into the mouse brain to establish electrical connection with the tissue. EOG responses were recorded at room temperature using an EPC 10 USB amplifier (HEKA Elektronik) and analyzed in PatchMaster Next (version 1.4.1). Signals were sampled at 10 kHz and low-pass filtered at 2.9 kHz. Response kinetics were evaluated by measuring the time to peak, defined as the interval from stimulus onset to the maximum response. All chemical reagents were obtained from Sigma-Aldrich.

### Odor-guided food-seeking test

Mice were tested for olfactory-guided food-seeking using a small piece of food buried beneath the cage bedding, ensuring that only olfactory cues were available. Both male and female mice (1–2 months old) were housed under a 12:12 h light/dark cycle and tested during the light phase. Animals were food-deprived overnight, with water provided ad libitum. Prior to testing, each mouse was familiarized with the experimental setup for 5 minutes.

For the test, the mouse was placed in a novel cage (30 cm × 12 cm × 15 cm) containing approximately 3 g of cookie (Oreo) buried under ∼2 cm of fresh bedding. The latency to locate the cookie was measured with a stopwatch, and successful retrieval was defined as digging it up with the forepaws, picking it up, and placing it in the mouth. A maximum search time of 5 minutes was allowed; if the cookie was not found within this period, the trial ended, the cookie was uncovered, and the mouse was allowed to eat it. The buried cookie’s position was randomized for each experiment. On the following day, the cookie was placed on top of the bedding. All mice, including those that did not locate the buried cookie within the allotted time, were included in the analysis.

### Psychophysical olfactory assessment

The study on the family was approved by the Ethics Committee of “Azienda Ospedaliero Universitaria Policlinico of Bari,” Italy (n.7343/2022).

We assessed olfactory function using the validated extended Sniffin’ Sticks test (SST) (Burghart Messtechnik, Holm, Germany), which includes tests for odor threshold (phenylethyl alcohol), odor discrimination, and odor identification. The maximum score for each of the 3 subsections of the SST is 16. Results from the 3 tests were presented as a composite TDI score (range 1–48) (Rumeau et al. 2016). To determine the olfactory abilities of our subjects we referred to (Hummel et al. 2007; Oleszkiewicz et al. 2019) where cut-off values are functional anosmia (TDI ≤ 16.5), functional hyposmia (16.5 < TDI <30.75), or functionally normal ability to smell (TDI ≥ 30.75) (Hummel et al. 2007 updated in Oleszkiewicz et al. 2019). Testing was conducted according to standard procedures described by (Hummel et al., 1997; 2007).

### Statistical analysis

Statistical analyses were conducted using Python 3.13.5 (Python Software Foundation, https://www.python.org/) in Visual Studio Code. Sample distributions were assessed for normality using scipy.stats.shapiro() (Virtanen et al., 2020). For parametric ANOVA, ols() and anova_lm() from Statsmodels (Seabold & Perktold, 2010) were used, followed by Tukey post hoc tests (pairwise_tukeyhsd()) (Benjamini & Hochberg, 1995). Alternatively, one-way and two-way ANOVA with post hoc pairwise comparisons were also performed using the Pingouin package (pingouin.anova(), pingouin.pairwise_ttests()) (Vallat, 2018), which allowed easy computation of effect sizes and p-value corrections. For nonparametric tests, scipy.stats.kruskal() and scipy.stats.mannwhitneyu() were used, with multiple testing corrections applied via multipletests(method=’fdr_bh’) (Benjamini & Hochberg, 1995). Linear mixed models (LMMs) were fitted using statsmodels.formula.api.mixedlm() (Seabold & Perktold, 2010). Data visualization was performed using Matplotlib (Hunter, 2007), NumPy (Harris et al., 2020), Pandas (McKinney, 2010), and itertools (Python Software Foundation, 2025), Matplotlib package, Inkscape computer software (Inkscape Project, 2022), Series of MALDI-TOF mass spectra (three replicates for each sample) were averaged using the software for the mass spectrometer ClinProTools 3.0 (Bruker Daltonics, Bremen, Germany) to find the area under the peaks. Statistically significant differences between series of mass spectra obtained from OE samples (WT, HET and MUT at P36 and P60) were determined using Anova when dependent variable was normally distributed and Kruskal-Wallis test when the distribution did not fit the normally distribution. A p-value < 0.05 was set as a threshold for statistically significant differences between peak intensity.

## Author Contributions

M.G.F. performed electrophysiological and behavioral experiments. M.G.F. and D.L. performed OE isolation. M.G.F., D.L. and R.P.G. performed immunofluorescence experiments. M.G.F., D.L. and R.P.G. performed image acquisition. S.L performed lipid analysis. M.G.F., S.L. and M.D. analyzed the experiments. M.D., M.G.F, D.L. and J.R., A.F. conceptualized and wrote the manuscript. All the authors approved the final version of the manuscript.

## Acknowledgments

We thank Leonardo Bombini and Rossella Vernò for technical support in the lab.

## Ethics Statement

All experiments were performed using procedures approved by the Institutional Committee on Animal Research and Ethics of the University of Bari and the Italian Health Department (Project n°465/2022-PR) and in accordance with the European directive on animal use for research. The study on the family was approved by the Ethics Committee of “Azienda Ospedaliero Universitaria Policlinico of Bari,” Italy (n.7343/2022).

## Data Availability Statement

The data that support the findings of this study are available from the first and corresponding authors upon reasonable request.

